# Expression of retrotransposons contributes to aging in *Drosophila*

**DOI:** 10.1101/2022.11.14.516438

**Authors:** Blair K. Schneider, Shixiang Sun, Moonsook Lee, Wenge Li, Nicholas Skvir, Nicola Neretti, Jan Vijg, Julie Secombe

## Abstract

Retrotransposons are a class of transposable elements capable of self-replication and insertion into new genomic locations. Across species, the mobilization of retrotransposons in somatic cells has been suggested to contribute to the cell and tissue functional decline that occurs during aging. Retrotransposon expression generally increases with age, and *de novo* insertions have been observed to occur during tumorigenesis. However, the extent to which new retrotransposon insertions occur during normal aging and their effect on cellular and animal function remains understudied. Here we use a single nucleus whole genome sequencing approach in *Drosophila* to directly test whether transposon insertions increase with age in somatic cells. Analyses of nuclei from thoraces and indirect flight muscles using a newly developed pipeline, Retrofind, revealed no significant increase in the number of transposon insertions with age. Despite this, reducing the expression of two different retrotransposons, *412* and *Roo,* extends lifespan, without increasing stress resistance. This suggests a key role for transposon expression and not insertion in regulating longevity. Transcriptomic analyses revealed similar changes to gene expression in *412* and *Roo* knockdown flies and highlighted potential changes to genes involved in proteolysis and immune function as potential contributors to the observed changes in longevity. Combined, our data show a clear link between retrotransposon expression and aging.

**Author Summary:** With the onset of modern medicine, the average age of the population has significantly increased, leading to more individuals living with chronic health issues. Rather than treat each age-associated disorder individually, one approach to target multiple health concerns simultaneously might to be target aging itself. Genomic instability is a hallmark of aging cells that has been proposed to be a key contributor to age-associated cellular decline. Transposons are mobile genetic elements capable of inserting into new genomic locations, thus having the potential to increase genomic instability. Consistent with this, transposon expression generally increases with age. However, the extent to which transposon insertions accumulate to disrupt the genome of cells within aging individuals has remained an open question. We specifically answer this through single cell whole genome sequencing and find that transposon insertions do not increase with age. Even though insertions did not increase, the expression of transposons is linked to aging, as reducing the expression of individual transposons extended lifespan. Transcriptome studies of these long-lived flies revealed increased expression of genes linked to proteolysis genes and to functioning of the immune system. Our study therefore establishes transposon expression, and not insertion, as a critical contributor to animal aging.

## Introduction

Across species, aging is associated with functional decline and an increased likelihood of one or more disorders that adversely affect quality of life (1, 2). Driving the changes that occur at the whole animal level are a range of alterations to cellular function. One hallmark of aging is genomic instability, in which the accumulation of mutations can alter critical gene expression programs and impact cell division by promoting tumor formation or by increasing cellular senescence (3, 4). One predicted genetic contributor to aging is the expression and mobilization of transposable elements (TEs) in somatic cells. TEs are abundant, comprising ~45-50% of the human, ~37% of the mouse, and ~20% of the *Drosophila* genomes (5–9). While the specific TEs found in each animal species are distinct, TEs are a universal feature of eukaryotic genomes whose expression generally increases with age (10–12). This increased expression relates to Class 1 transposons, also known as retrotransposons (RTs), that replicate using an RNA intermediate or a ‘copy, paste’ mechanism (12). RTs are therefore able to increase their genomic copy number over time, which gives them a high mutagenic potential. In contrast, class 2 transposons (DNA transposons) mobilize by excising themselves in a ‘cut, paste’ mechanism, so their total number does not increase over time (12) and less is known about their expression with respect to aging. However, the precise role that TEs play in driving age-related phenotypes remains an open question.

The most obvious consequence of TE expression with age is the potential to cause genomic instability through insertional mutagenesis and/or the creation of insertions/deletions because of the double-stranded DNA breaks that are needed for TE reinsertion (12, 13). In addition, cDNA generated as an intermediate during RT mobilization can activate the immune response and lead to chronic inflammation in mammals (14–16). Consistent with their potential for interfering with cellular function, several mechanisms that are conserved across species have evolved to limit the expression of transposons. For example, many transposons in the genome are contained within constitutive heterochromatin, which is largely transcriptionally silent (3, 9, 17–20). For those elements inserted within euchromatin, RT expression is regulated post transcriptionally via the siRNA pathway that degrades double-stranded RNA complexes (21, 22). Further supporting a role for heterochromatin and RNAi in repressing TE’s influence on aging, modulating the activity of either of these pathways can alter lifespan. For example, reducing the expression of the heterochromatin component *Lamin B* or components of RNAi-mediated TE silencing machinery decreases lifespan in *Drosophila* (11, 20). Conversely, increasing expression of the heterochromatin-promoting histone methyltransferase, *su(var)3-9,* or the activity of the RNAi pathway increases lifespan (20, 23, 24). It is, however, notable that heterochromatin and the RNAi pathway are not specific to the repression of TEs, which complicates the interpretation of the changes to lifespan observed. More direct evidence supporting a link between TE activation and aging has come from the use of reverse transcriptase inhibitor drugs that broadly inhibit the ability of RTs to replicate. For example, *Drosophila* fed nucleoside reverse transcriptase inhibitors (NRTIs) have an increased lifespan compared to controls (25). This is also observed in mice, where NRTI treatment attenuates the shortened lifespan caused by loss of *SIRT6,* a known repressor of *LINE1 (L1)* elements (14).

The key to understanding the link between TEs and aging is the extent to which their expression leads to new insertions. Analyses of the TE, *mdg4* (formerly known as *Gypsy*), in the adult *Drosophila* brain and fat body using a reporter revealed an increase in the number of insertions with age (20, 23, 26). However, expanding these findings to endogenous TEs has been challenging, as each somatic cell contains a unique set of insertions which are difficult to detect using bulk whole genome sequencing approaches. One approach to address this has been to develop bioinformatic tools to detect TE insertions more accurately, for example, by using TE junction and target site duplication data. When applied to bulk whole genome sequencing (WGS), this approach can detect an age-associated increase in TE insertion number in fly strains with reduced RNAi pathway activity and in clonally expanded tumors, but not in wild-type animals (24, 27). New insertions can also be accurately identified using long-read sequencing of bulk DNA as whole elements are detected rather than breakpoints (24, 27). While a small number of *de novo* insertions were observed using this technique, it was difficult to determine how frequently TEs mobilized. While technologies to detect new TE insertions in somatic cells have become more robust using bulk sequencing approaches, the frequency of new TE insertions within individual somatic cells during aging remains unknown.

To understand the extent to which TEs mobilized during aging, we took a single nucleus whole genome sequencing (WGS) approach using a new pipeline called Retrofind, that allows us to accurately define the insertional position and load per cell. Using nuclei isolated from adult thoraces or indirect flight muscles (IFMs), we found that the number of TE insertions does not increase with age. However, reducing the expression of two individual TEs, *412* and *Roo,* led to lifespan extension. This suggests a key role for the expression, and not insertion, of TEs impacting lifespan. This increased lifespan did not correlate with improved stress resistance or other health improvements traditionally associated with longer life. Transcriptomic studies of long-lived TE knockdown flies revealed that the expression of genes involved in proteolysis were upregulated, including the *Jonah* family of genes that encode serine hydrolases. Additionally, antimicrobial peptides (AMPs), the downstream products of activation of the *Drosophila* innate immune system that is similar to mammalian inflammation, were dysregulated in knockdown animals. Overall, our studies show that TE expression and not insertion likely contributes to aging, potentially through regulation of the *Jonah* genes and immunity.

## Results

### TE insertions do not increase with age

To identify *de novo* TE insertions, we used a single nucleus whole genome sequencing approach (Fig 1A). Using this methodology, new insertions are represented in approximately half of the reads in each nucleus, facilitating robust detection. We carried out our studies using the well characterized *w^1118^* strain that shows a typical lifespan and is often used a wild-type control strain (Fig 1A)(24, 28). To distinguish new insertions that occur with age from preexisting ones within the germline genome, we defined the TE landscape in the *w^1118^* strain by carrying out bulk sequencing from pooled young thoraces to a total depth of 157x coverage. Sequencing data was analyzed using a newly developed in-house TE detection pipeline called RetroFind, in addition to a published pipeline that has successfully detected somatic insertions in clonally expanded tumors, so serves as independent validation (27). New TE insertions were identified by similar criteria for each pipeline by detecting both split and discordant reads of evidence from paired-end sequencing data. Additionally, newly called insertion sites in the germline genome needed to possess the target site duplication that occurs because of the double stranded breaks made by the TE-encoded integrase. A total of 871 TE insertions unique to *w^1118^* relative to the *dm6* reference genome were detected by the two pipelines, with 505 being detected by both (Fig 1B, S1 Table). We consider these to be high confidence insertions. TE insertions identified in *w^1118^* were primarily within intronic and intergenic regions and were largely excluded from promoters (+/- 100 base pairs of the transcription start site) and coding sequences (CDS) where they might disrupt gene function (Fig 1C).

**Fig 1.**
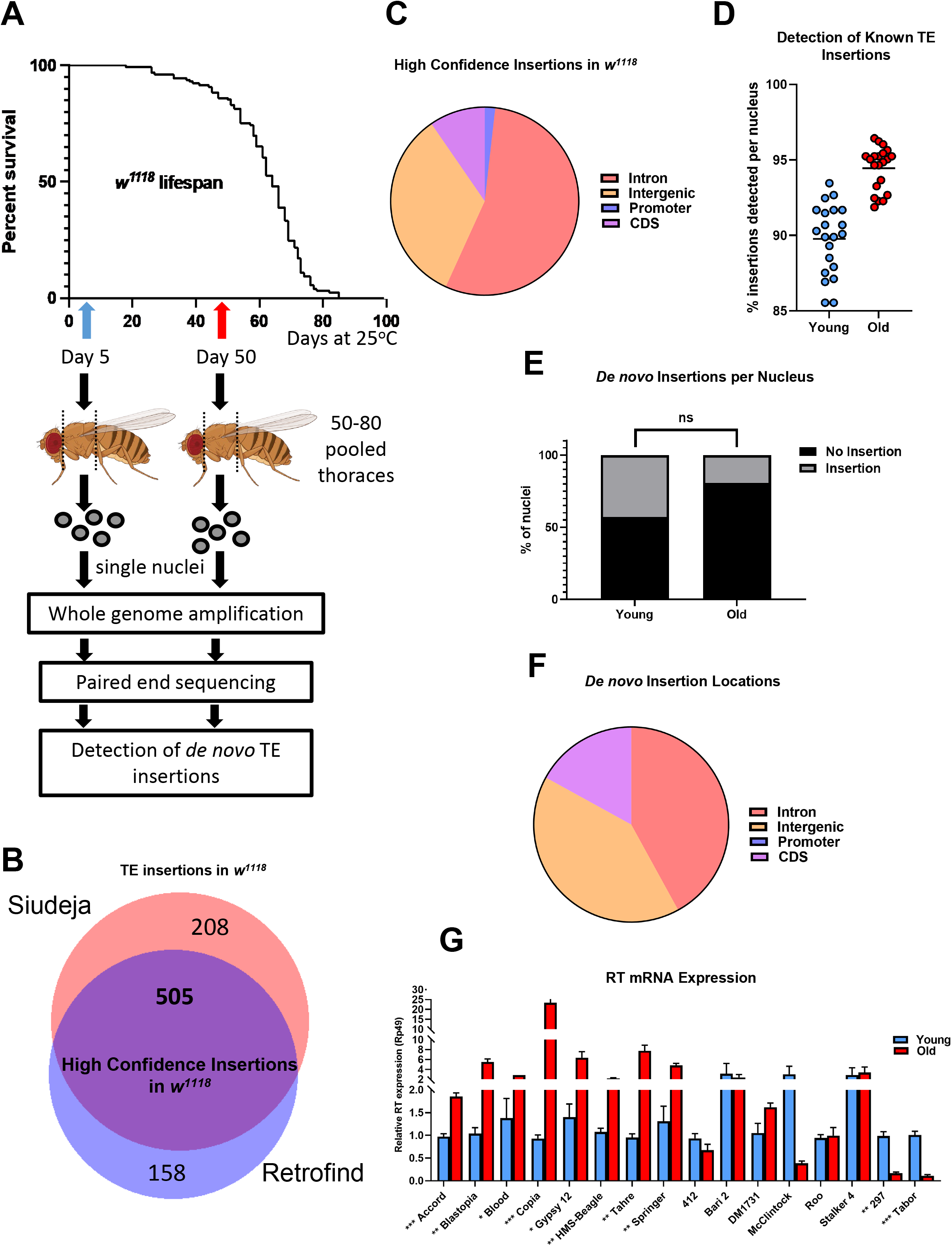
Single cell whole genome sequencing of young (day 5) and old (day 50) *Drosophila* thoraces. (A) A schematic of the workflow of the isolation of single nuclei and how new TE insertions were detected. The typical *w^1118^* wild type strain lifespan is also displayed. (B) The number of new TE insertions in the *w^1118^* strain from the bulk WGS in comparison to the sequenced reference strain (dm6) using both the pipeline from Siudeja et al (27) and Retrofind. 505 insertions were called by both pipelines. (C) The distribution of the genomic locations where the *w^1118^* strain TE insertions fall. (D) The number of known TE insertions (strain/background insertions) that were able to be detected within each nuclear sample represented as a percentage of the insertions detected in the sample out of total strain insertions. Each dot represents a nucleus. (E) The number of new TE insertions within each nucleus that are not previously called within the strain or other nuclei. Each dot represents a nucleus. Unpaired t-test. ns p= 0.1809. (F) The distribution of genomic locations where the 15 new TE insertions within individual nuclei fall. (G) qPCR using SYBR green showing levels of TE mRNA relative to Rp49 (Rpl32) from adult *w^1118^* flies young (day 5) in blue and old (day 50) in red head and thoraces. The experiment was performed in three biological replicates. An unpaired t-test with Bonferroni correction was used. *** *(Accord)* p= 0.0009 ** *(Blastopia)* p= 0.0020. * (*Blood*) p= 0.0262. *** *(Copia)* p= 0.0007. * *(gypsy12)* p= 0.0171. ** *(HMS-Beagle)* p= 0.0011. ** *(Tahre)* p= 0.0038. ** *(Springer)* p= 0.0026. ns (*412*) p= 0.1453. ns *(Bari2)* p=0.7403. ns *(DM1731)* p= 0.0702. ns *(McClintock)* p= 0.1814. ns (*Roo*) p= 0.7985. ns *(Stalker4)* p= 0.7957. ** (*297*) p= 0.0012. *** *(Tabor)* p= 0.0005.

To determine if the number of TE insertions increased with age in somatic cells, we isolated and amplified genomic DNA from 21 individual nuclei from thoraces of young (5 days old) and from old (50 days old) flies (Fig 1A). Whole genome sequencing (WGS) of these 42 nuclei revealed an average 317x coverage of the genome for nuclei from young thoraces and 354x for nuclei from old thoraces. To confirm our ability to map TEs in individual nuclei, we looked for the 505 high confidence TE insertions detected from sequencing bulk genomic DNA. Confirming a robust ability to detect TEs, we find an average of 90% of high confidence insertions in nuclei from young animals and 94% from old animals (Fig 1D). While it is notable that more of the known *w^1118^* insertions were detected in nuclei from old animals, this likely reflects the higher sequencing depth observed from these samples.

To identify *de novo* insertions, we used Retrofind and the published pipeline to analyze the sequencing data for each nucleus, with unique insertions detected by both pipelines being deemed high confidence (27). This revealed that 57% and 76% of nuclei from young and old animals, respectively, did not have any new TE insertions (Fig 1E). A total of 15 new insertions were found across 9 nuclei from young animals and 4 from old. A maximum of two insertions was detected in any individual nucleus. Each of the 15 *de novo* TE insertions detected was visually confirmed using Integrated Genome Viewer (IGV), similar to previous studies (27). Like existing TE insertions within the genome, new insertions occurred primarily in non-coding regions of the genome (Fig 1F). Most of the new insertions were observed in nuclei from young flies, and all the insertions observed in old flies, were the *hobo* element (also known as the *H-element)* (S2 Table). This DNA terminal inverted repeat transposon is an evolutionarily recent addition to the *Drosophila* genome that is known to be active (29). In addition, the long terminal repeat (LTR) retrotransposon *HMS-Beagle* and the DNA *S terminal inverted repeat element* showed a new single insertion in the somatic genome, each in a single nucleus from young animals. Based on these data, we tested whether the *w^1118^* strain shows increased TE expression with age. Using qPCR, we find that the expression of a subset of TEs increases expression with age (Fig 1G). For instance, the TEs *Gypsy12* and *Copia* that have been previously shown to increase their expression with age in other tissues such as the fat body (11). The expression of other elements did not change with age, such as *412* and *Roo*, and some showed decreased expression, such as *297* (Fig 1G). Thus, increased expression of TEs with age does correlate with additional insertional events.

Because our single nuclei were isolated from thoraces, they likely represent several different cell types, possibly obscuring an increase in TEs that might be observed by analyzing a single tissue. We therefore also purified nuclei from indirect flight muscles (IFMs), which are a major muscle group in the thorax that show age-associated decline (30–33). To purify IFM nuclei, they were labeled by a nuclear membrane localized GFP (UAS-GFP:KASH) using UH3-Gal4 (UH3>GFP:KASH; Fig 2A) (34). Individual GFP positive nuclei from dissected thoraces were isolated by fluorescence-activated cell sorting and whole genome amplified in a similar manner to our analyses of thoracic nuclei. Unamplified, bulk DNA from the abdomen and head regions of the same flies were sequenced to a total of 443x coverage to exclude strain-specific TE insertions. Sequencing of the individual nuclei revealed an average of 64x coverage from young nuclei and 115x from old. Like our findings using nuclei from thoraces, analyses of six and seven IFM nuclei from young and old flies, respectively, revealed no significant increase in TE insertions with age (Fig 2B). Approximately half of nuclei examined had no new insertions and the small number that were observed were inserted into intergenic or intronic sequences (Fig 2C). These data show that TE insertions do not increase significantly during aging in cells of the thorax or IFMs.

**Fig 2.**
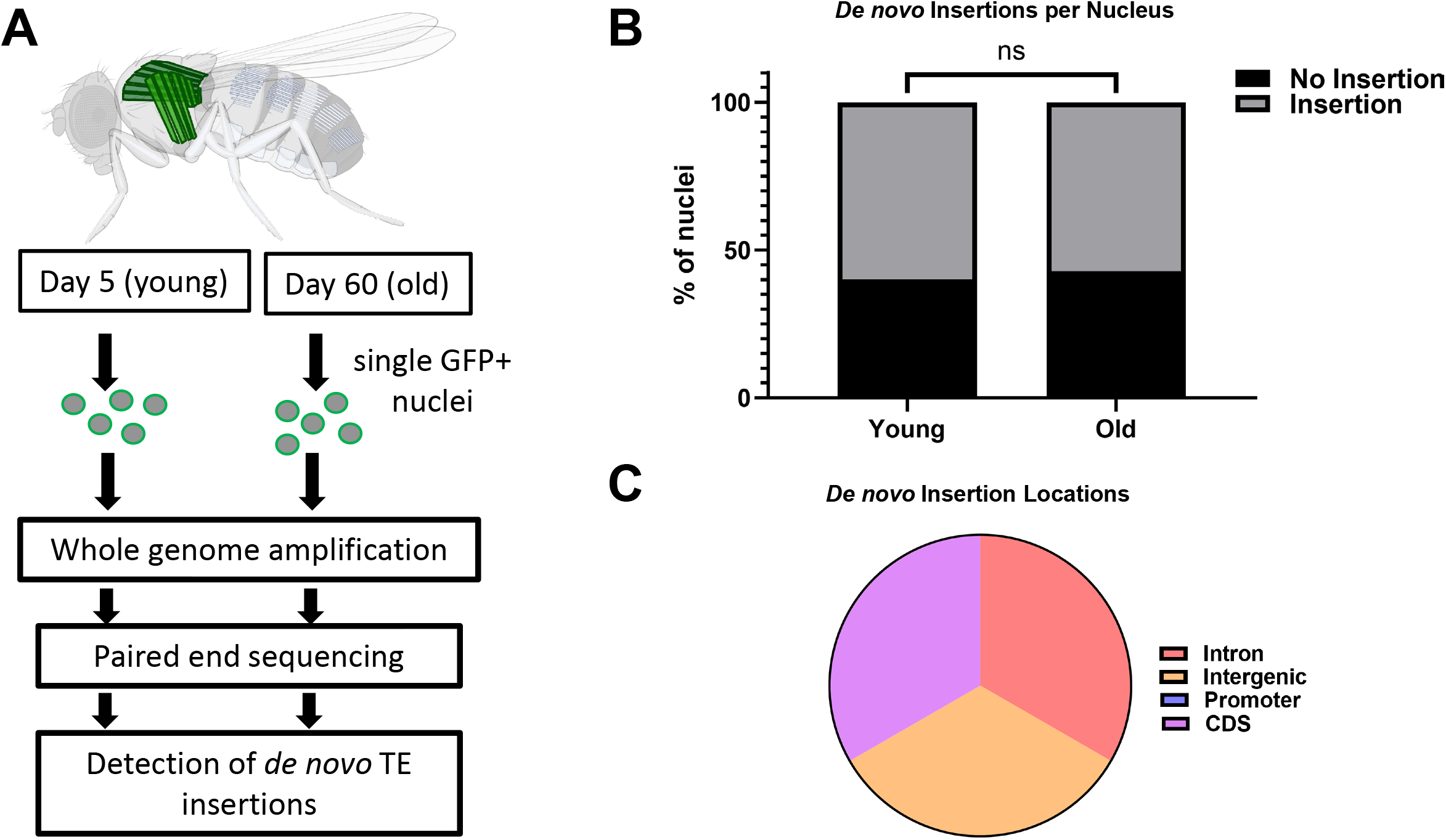
Single cell whole genome sequencing of young (day 5) and old (day 60) *Drosophila* indirect flight muscles. (A) A schematic of the workflow of the isolation of single nuclei and how new TE insertions were detected from indirect flight muscles (IFMs). (B) The number of new TE insertions within each nucleus that are not previously called within the strain or other nuclei. Each dot represents a nucleus. Unpaired t-test. ns p >0.9999. (C) The distribution of genomic locations where the new TE insertions within individual nuclei fall.

### Reduced expression of individual retrotransposons extends lifespan

While no increase in TE insertions were observed during adulthood, the expression of these elements could impact cell function, thereby altering lifespan. For this reason, we examined the effect of attenuating retrotransposon expression. We chose to focus initially on the retrotransposon *412*, which is part of the *gypsy* super-family of LTR elements that has been previously shown to be stable in the germline (35). Consistent with this, we did not observe any new *412* insertions in *w^1118^* compared to the sequenced strain (S1 Table). Like most RTs, *412* is multi-copy within the genome, having 36 copies in total, 24 of which are full length (36). We therefore used a knockdown approach to reduce the expression of *412* to assess the effect on life- and healthspan. To do this, we generated two UAS-regulated short hairpin transgenes, 412#1, and 412#2, in addition to a control construct expressing a shRNA predicted not target any mRNAs (control). Driving the expression of UAS-shRNA transgenes targeting *412* with the ubiquitous Actin5C-Gal4 (Act5C>shRNA) driver led to a ~2-fold reduction in *412* mRNA levels (Fig 3A). *412* knockdown flies completed metamorphosis normally and were grossly morphology normal (S1 Fig). Quantifying the lifespan of these animals revealed significantly extended median and maximum lifespan compared to controls (Fig 3B-D).

**Fig 3.**
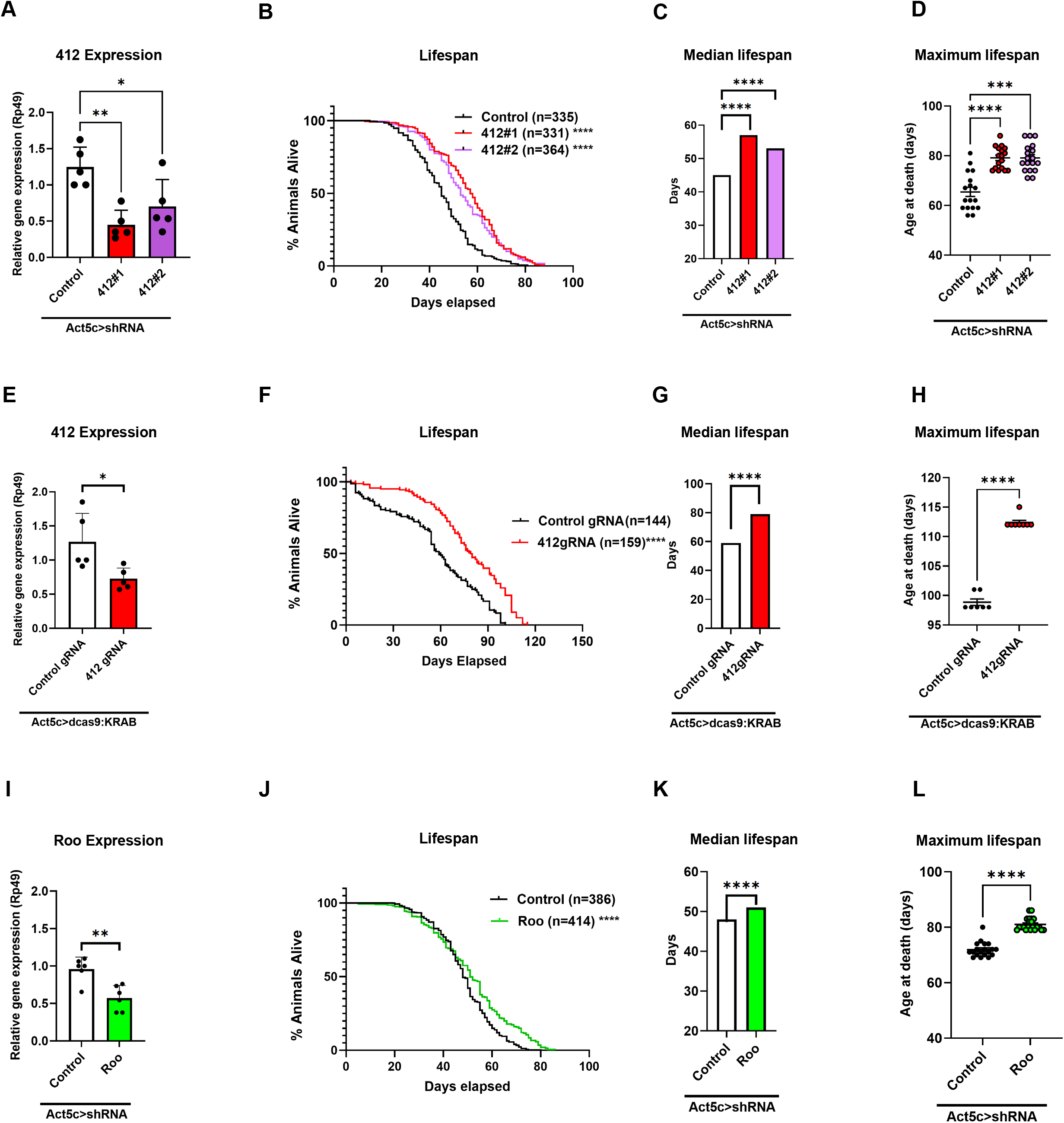
Lifespan of ubiquitous shRNA and CRISPRi knockdown of the retrotransposons, *412* and *Roo*. (A) qPCR using SYBR green showing levels of *412* mRNA relative to *Rp49 (Rpl32)* from adult flies expressing a control shRNA transgene under the control of Act5C-Gal4 compared to the *412* shRNA knockdowns (Act5C>shRNA). The experiment was performed in five biological replicates. An unpaired t-test with Bonferroni correction was used. ** p=0.0018. * p=0.0219. (B) Survival of the *412* shRNA lines compared to control shRNA driven ubiquitously by Act5C (Act>shRNA). Dunnett test. ****p <0.0001. (C) Median lifespan of the *412* shRNAs compared to control shRNA driven ubiquitously by Act5C (Act>shRNA). Gehan-Breslow-Wilcoxon test. ****p<0.0001. (D) Maximum lifespan of the *412* shRNAs compared to control shRNA driven ubiquitously by Act5C (Act>shRNA). Permutation test (95th percentile/top 5%) followed by two-sided t-test with correction for multiple comparisons. ****p<0.0001. ***p= 0.0001. (E) qPCR using SYBR green of 412gRNA CRISPRi compared to the control gRNA control (Act5C>dcas9-KRAB) relative to *Rp49* (*Rpl32*). An unpaired t-test with Bonferroni correction was used. * p=0.0269. (F) Survival of the CRISPRi driven ubiquitously by Act5C (Act>dcas9KRAB) with the 412gRNA compared to control gRNA. Dunnett test. **** p <0.0001. (G) Median lifespan of the CRISPRi driven ubiquitously by Act5C (Act>dcas9KRAB) with the 412gRNA compared to control gRNA. Gehan-Breslow-Wilcoxon test. **** p <0.0001. (H) Maximum lifespan of the of the CRISPRi driven ubiquitously by Act5C (Act>dcas9KRAB) with the 412gRNA compared to control gRNA. Permutation test (95th percentile) followed by two-sided t-test with correction for multiple comparisons. * p=0.0377. (I) qPCR using SYBR green of the *Roo* mRNA KD compared to the control shRNA control (Act5C>shRNA) relative to *Rp49* (*Rpl32*). The experiment was performed in six biological replicates. An unpaired t-test with Bonferroni correction was used. ** p=0.0022. (J) Survival of the *Roo* shRNA compared to control shRNA driven ubiquitously by Act5C (Act>shRNA). Dunnett test. **** p <0.0001 (K) Median lifespan of the *Roo* shRNA compared to control shRNA driven ubiquitously by Act5C (Act>shRNA). Gehan-Breslow-Wilcoxon test. **** p <0.0001. (L) Maximum lifespan of the *Roo* shRNA compared to control shRNA driven ubiquitously by Act5C (Act>shRNA). Permutation test (95th percentile) followed by two-sided t-test with correction for multiple comparisons. **** p<0.0001.

To confirm the lifespan extension caused by RNAi-mediated knockdown of *412*, we additionally used CRISPR interference (CRISPRi) to reduce expression of this TE. To do this, we generated a UAS transgene encoding an enzymatically dead Cas9 protein fused to the *KRAB* transcriptional repressor (UAS-dCas9:KRAB). dCas9:KRAB was targeted to the *412* LTR enhancer/promoter region using a transgene expressing a guide RNA (gRNA) under the control of the ubiquitously expressed, *U6* promoter (412gRNA). As a control, we generated control non-targeted gRNA (control gRNA). CRISPRi was carried out by crossing Act5C>dCas9:KRAB and 412gRNA flies, which led to a 1.7-fold reduction in 412 mRNA levels (Fig 3E). In contrast to 412 knockdown adult flies that were indistinguishable from controls, 412gRNA CRISPRi flies were 7% heavier than control gRNA flies although they were morphologically normal (S1 Fig). Mirroring our shRNA results, 412 gRNA-expressing flies showed extended median and maximum lifespan compared to control gRNA-expressing flies (Fig 3F-H). Reducing the expression of a single RT is therefore sufficient to extend lifespan.

To test whether changes in lifespan were specific to *412*, we reduced the expression of another RT that showed 112 new insertions in *w^1118^* compared to the annotated *Drosophila* genome (S1 Table). Driving the expression of a UAS-shRNA transgene targeting *Roo* using Act5C-Gal4 led to a ~2-fold reduction in mRNA levels and did not adversely affect ability of animals to complete metamorphosis or their gross morphology (Fig 3I; S1 Fig). Quantifying the lifespan of these animals revealed a significantly extended median and maximum lifespan compared to control animals (Fig 3J-L). The expression of *412* and *Roo* therefore both contribute to aging.

### Long-lived 412 or Roo knockdown flies do not show improved locomotion or stress resistance

Extension of longevity can be associated with improved healthspan, which can be measured as a delay in the onset of age-associated phenotypes. For example, long-lived fly strains, such as those with reduced insulin signaling, show increased resistance to starvation and oxidative stress (37). Additionally, progeroid flies have an accelerated decline in locomotor activity with age (38). We therefore tested the extent to which flies with reduced expression of *412* or *Roo* showed improvements in locomotion and/or resistance to a range of stress conditions. One classic indicator of age-associated decline is a reduced locomotion, which can be quantified using a negative geotaxis assay. This assay takes advantage of an innate response whereby flies move against gravity by climbing to the top of a vial after being tapped to the bottom (39). Young, healthy, flies quickly climb to the top third of the vial while fewer older flies climb this distance. As expected, locomotor ability declined with age across genotypes, with those in midlife (18 days old) and old age (40 days old) showing an attenuated negative geotaxis response than young flies (5 days old) (S2 Fig). Midlife flies also have more locomotion when compared to old age flies (S2 Fig). At all ages tested, *412* knockdown flies displayed locomotor capacity that was either indistinguishable or worse (412#1) than control animals (412#2) (Fig 4A and S2 Fig). Similarly, the locomotor capacity of *Roo* knockdown animals was not different than control animals (Fig 4A).

**Fig 4.**
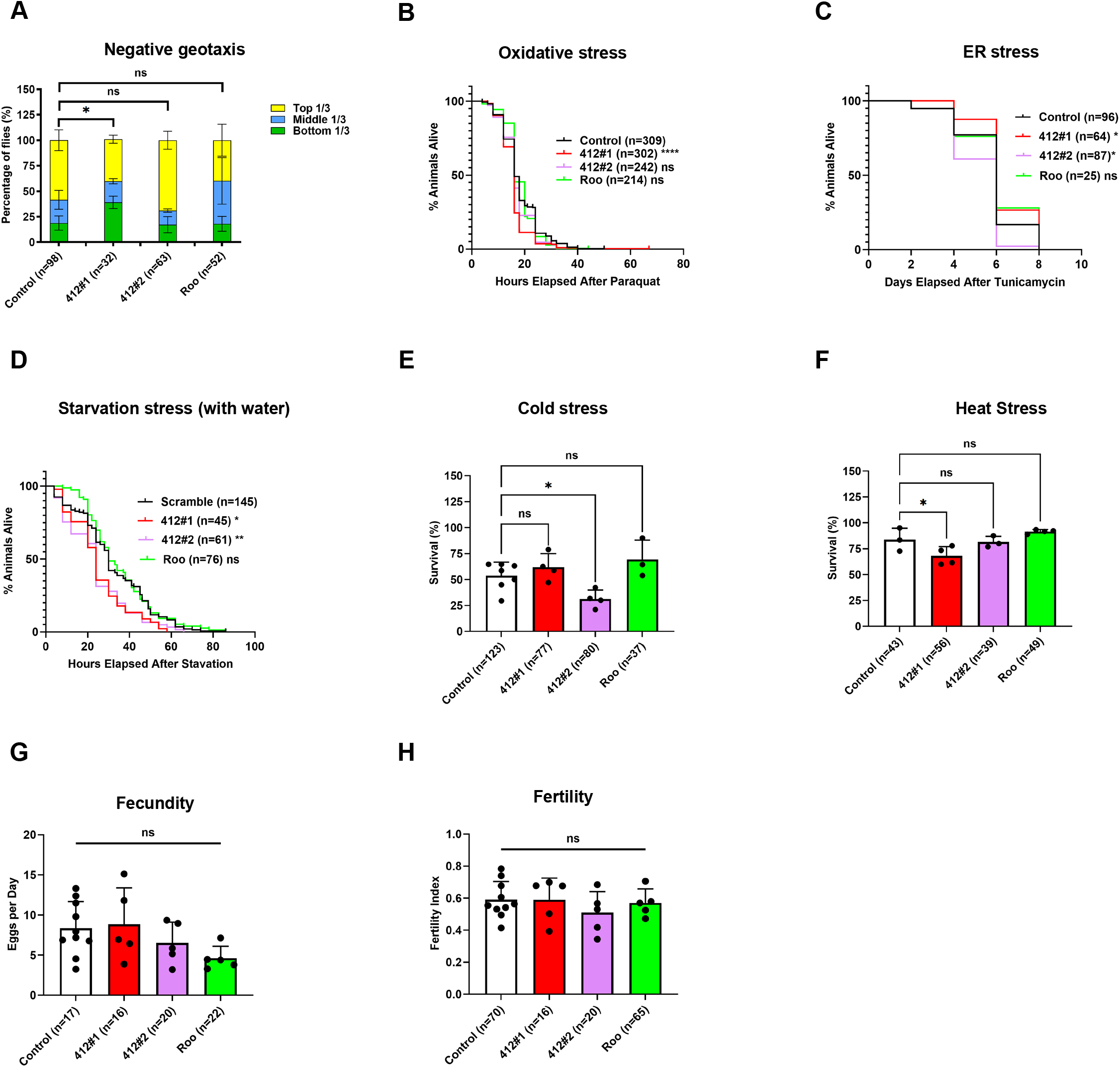
Stress resistance assays of *412* and *Roo* knockdown animals. (A) Act5c>shRNA day 18 measurement of locomotion via negative geotaxis assay. The percentages of flies in each third of the vial is displayed. Chi-square test for trend. * (412#1) p=0.0428. ns (412#2) p=0.1434. ns (Roo) p=0.1934. (B) Act5c>shRNA day 40 response to oxidative stress by feeding paraquat and measuring survival. Dunnett test. **** (412#1) p<0.0001. ns (412#2) p= 0.0702. ns (Roo) p >0.9999. (C) Act5c>shRNA day 20 response to endoplasmic reticulum (ER) stress by feeding tunicamycin and measuring survival. Dunnett test. ns (412#1) p=0.0826. ** (412#2) p=0.0046. ns (Roo) p=0.1721. (D) Act5c>shRNA day 40 response to starvation by only giving the flies access to water and measuring survival. Dunnett test. ns (412#1) p=0.8332. ns (412#2) p=0.5694. ns (Roo) p=0.2239. (E) Act5c>shRNA day 20 response to cold stress by keeping flies at 4 degrees Celsius and measuring survival after 48-hours recovery. Each dot is a vial or replicate of approximately 20 flies. Fisher’s exact test. ns (412#1) p=0.0703. * (412#2) p= 0.0461. ns (Roo) p= 0.6592. (F) Act5c>shRNA day 20 response to heat stress by keeping flies at 37 degrees Celsius and measuring survival after 48-hours recovery. Each dot is a replicate of approximately 20 flies. Fisher’s exact test. ns (412#1) p= 0.0915. ns (412#2) p= 0.7643. ns (Roo) p= 0.5061. (G) Act5c>shRNA day 25 measurement of fecundity. Data is displayed as average number of eggs laid per day over 5 days. Each dot represents a day for the number of flies in one vial. One-way ANOVA. ns (412#1) p=0.9925. ns (412#2) p=0.7231. ns (Roo) p=0.1803. (H) Act5c>shRNA day 25 measurement of fertility. Fertility index is calculated as number of progeny divided by number of eggs laid. Data is displayed as average fertility index over 5 days. Each dot represents a day for the number of flies listed. One-way ANOVA. ns (412#1) p>0.9999. ns (412#2) p=0.5881. ns (Roo) p=0.9887.

To assess whether reduced RT expression altered resistance to oxidative stress, we treated ubiquitous *412* or *Roo* knock down flies with paraquat and quantified survival compared to control flies. This revealed that the two *412* shRNA transgenes behaved differently from each other, with 412#2 showing no change to survival and 412#1 having slightly reduced resistance to oxidative stress (Fig 4B). *Roo* knockdown animals showed no change resistance to oxidative stress compared to control animals (Fig 4B). We additionally tested survival in response to endoplasmic reticulum (ER) stress through treatment with the antibiotic tunicamycin (40), starvation, and thermal stress (cold and heat). None of these treatments led to a consistent change in survival for *412* or *Roo* knockdown flies except starvation, where both *412* knockdowns showed increased sensitivity (Fig4C-F). An additional corollary to increased lifespan is a decline in fertility (41). We therefore quantified fecundity of *412* and *Roo* knockdown flies by counting the number of eggs laid per day and found no significant difference (Fig 4G). Nor was the fertility of *412* or *Roo* knockdown animals altered, as the number of adult flies produced from those eggs was indistinguishable from controls (Fig 4H). Combined, these assays show that reducing *412* or *Roo* expression does not improve standard assays of health and/or stress resistance that might be expected in these long-lived flies.

### TE knockdown affects genes linked to proteolysis and immunity

To gain insight into the cellular changes caused by reduced TE expression, we performed RNA-seq on thoraces of Act>shRNA animals at mid-life (day 20). We chose mid-life as others have seen changes to age-associated phenotypes at this time point (20, 38). 332 differentially expressed genes (DEGs) were identified in *412* knockdown animals compared to control shRNA expressing flies using a 5% false discovery rate (FDR) cutoff. 212 of these genes were upregulated while 120 were downregulated and averaged a log2 fold change of 2.5 and 1.5, respectively (Fig 5A; S3 Table). Functional analyses of upregulated genes using GO DAVID revealed significant enrichment in the single gene ontology (GO) category of proteolysis using a 1% FDR (42, 43). Many of these genes were members of the Jonah (Jon) family of serine proteases, including *Jonah 25Bi-iii, Jonah 44E, Jonah 65Aiii, Jonah 65Aiv, Jonah 74E,* and *Jonah 99Ci-iii* (S5 Table) (42, 43). Similar GO analyses of downregulated genes did not reveal any significantly enriched categories using an FDR of 1%.

**Fig 5.**
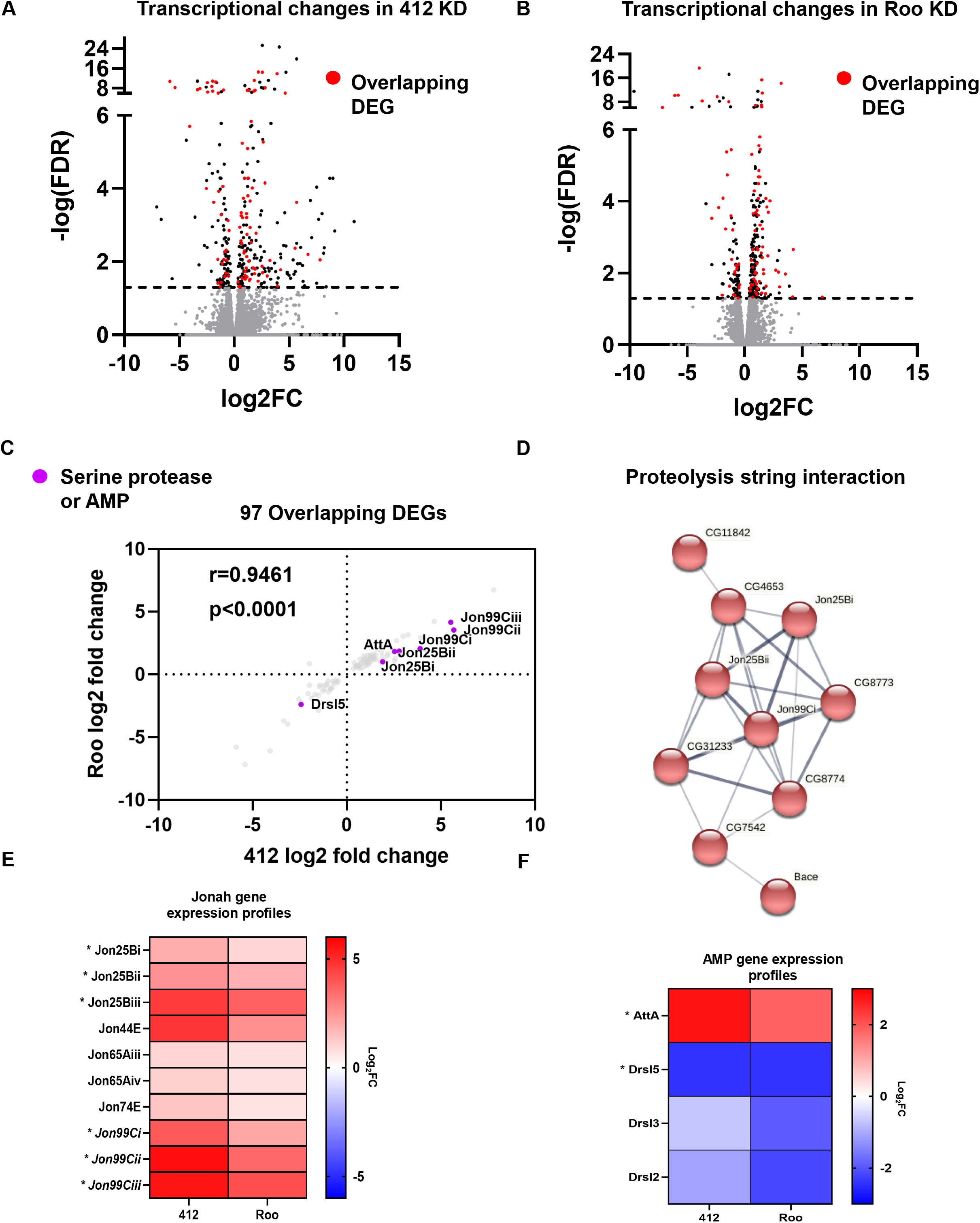
Transcriptomic analysis of *412* and *Roo* knockdown animals. (A) Volcano plot of differentially expressed genes (DEGs) from whole thorax of Act5c>412#1 shRNA knockdown animals compared to control shRNA animals. Genes with a false discovery rate (FDR) <0.05 are highlighted in black and above the dashed line. Genes that are differentially expressed in both *412* and *Roo* knockdown animals (overlapping genes) are highlighted in red. (B) Volcano plot of differentially expressed genes (DEGs) from whole thorax of Act5c>Roo shRNA knockdown animals compared to control shRNA animals. Genes with a false discovery rate (FDR) <0.05 are highlighted in black and above the dashed line. Genes that are differentially expressed in both *412* and *Roo* knockdown animals (overlapping genes) are highlighted in red. (C) Correlation of Log2FoldChange (Log2FC) of the overlapping DEGs between *412* and *Roo* knockdown animals. r=0.9461. Deming regression. p<0.0001. Serine proteases and AMPs that are also DEGS are enlarged and highlighted in purple. (D) Protein clustering performed on the overlapping DEGs using String with single nodes removed and 1 cluster used. There is a serine protease cluster of genes interacting. (E) Heatmap of log2fold change (Log2FC) from RNA-seq data across samples of the *Jonah (Jon)* genes. The * marked genes are significantly differentially expressed in both *412* and *Roo* knockdown thoraces. All other genes are significantly differentially expressed in just *412* knockdown thoraces. (F) Heatmap of log2fold change (Log2FC) across samples of the antimicrobial peptides (AMPs). The * marked genes are significantly differentially expressed in both *412* and *Roo* knockdown thoraces. All other genes are significantly differentially expressed in just *412* knockdown thoraces.

To determine the extent to which knockdown of *Roo* led to similar changes to gene expression as *412*, we carried out RNA-seq analyses using thoraces of ubiquitous *Roo* knockdown flies. 344 genes were found to be differentially expressed in *Roo* knockdown animals, 236 of which were upregulated average and 108 were down compared to control animals (5% FDR; Fig 5B; S4 Table). As with knockdown of *412*, the changes to gene expression in response to reduced *Roo* expression were relatively small, averaging 1.1 and 1.3 log2 fold change for up- and downregulated genes, respectively. Interestingly, upregulated genes were enriched for the same proteolysis GO term as was observed in *412* knockdown animals (42, 43) (1% FDR cutoff; S6 Table). No GO terms were significantly enriched among the downregulated genes.

Based on the identification of the same GO term in *412* and *Roo* knockdown datasets, we compared the transcriptional changes of these two strains. This revealed 97 genes common to both datasets, a majority of which behaved similarly (r=0.9461; p<0.0001; Fig 5C). These genes were enriched for the single GO term of proteolysis (42, 43) (1% FDR cutoff; S7 Table). To understand the relationship between genes observed to be dysregulated within the proteolysis GO category, we used STRING, which revealed a distinct interaction node mainly based on co-expression and co-occurrence (Fig 5D) (44). The *Jon* genes, a family of serine proteases, were at the center of this node (45). Two thirds of the genes affected by *412* and *Roo* knockdown were upregulated in knockdown flies, including all of the genes within the proteolysis category (Fig 5E). While little is known about the Jonah proteins, their expression appears to be primarily in the gut, where they are assumed to aid in digestion (45). However, expression of the *Jon* genes has also been linked with changes to the immune deficient (IMD) and Toll immunity pathways, which have previously been associated with lifespan regulation (46–48). Consistent with this, the expression of antimicrobial peptides (AMPs) that are downstream of the IMD and Toll pathways are affected in *412* or *Roo* knockdown flies, with several *Drosomycins* (e.g., *Drsl2, Drsl3, Drsl5)* significantly downregulated and *Attacin-A (AttA)* being significantly upregulated (Fig 5F). Overall, our data suggest that TEs such as *412* and *Roo* alter the expression of genes related to proteolysis regulation and the immune system to influence longevity.

## Discussion

In this study we find that there is no significant increase in TE mobilization with age in *Drosophila,* indicating that they have a robust mechanism for preventing *de novo* TE insertions in somatic cells. Based on the observation that reduced activity of the RNAi pathway leads to an increase in TE insertions with age, this mechanism is likely a key means of limiting *de novo* TE insertions during normal aging (24). By combining a single nucleus whole genome sequencing approach with analyses using two pipelines, we are confident that new insertions would have been detected if present. Prior to our study, the most compelling data indicating increased TE insertions with age in wild-type animals came from use of a TE reporter transgene and sequencing of bulk DNA samples (24, 27, 49). Use of a fluorescent reporter for the RT *mdg4* revealed age-associated *de novo* insertions in cells of the adult brain and fat body, although the frequency was low (23, 26). In addition, examining the insertional proficiency of all TEs through long-read sequencing of pooled adult brains or midguts showed new insertions by several different elements, including *mdg4* and *Roo* (27). Like the *mdg4* reporter data, the number of new insertions detected using this approach was low, averaging less than one new integration event per individual (27). Because both approaches identified new insertions, it is possible that TEs are more highly expressed in the gut and brain than cells of the thorax, allowing some insertions to occur during aging in these tissues. Alternatively, given the low frequency of new TE insertions that were observed, it is more likely that these published data are congruent with our study. Based on the rate of transposition observed, our sequencing analyses of 42 thoracic nuclei (21 from young flies and 21 from old) was unlikely to be sufficient to detect new insertions. Consistent with our data that endogenous expression of TEs does not necessary lead to new insertions, overexpression of *mdg4* does not increase the number of genomic copies of this element (50). We therefore suggest that new TE insertions are unlikely to be a key driver of cell and tissue dysfunction that occurs during normal aging. Interestingly, this contrasts with disorders such as cancer where there is clear evidence from mammalian cells and flies that TE insertions are a frequent occurrence that likely impact disease severity (27, 51–53).

Previous functional evidence supporting TE mobilization playing a role in aging has come from the pharmaceutical approach of using RT inhibitors, with phosphonoformic acid (PFA) or dideoxyinosine (ddI) and Lamivudine (3TC) or Stavudine (d4T) treatment extending lifespan in *Drosophila* and mice, respectively (14, 25). NRTIs are well-established to block retrotransposon replication thereby preventing RT reinsertion (14). However, RT inhibitor treatment can additionally decrease RNA levels of the *L1* element in human cells, thus the effect of these drugs may not be limited to restricting mobilization (49). If NTRIs exert similar effects in flies, the extended lifespan seen using these drugs could be due to reduced expression of some or all TEs, rather than changes to replicative capacity. These data, like ours showing that TE knockdown affects lifespan without increasing mobilization, suggest that the presence of TE mRNA may be generally detrimental to cell and animal function.

Because of their similarity to retroviruses, cytosolic DNA intermediates produced by RTs can trigger the activity of the immune system (14–16, 54). In human cells, *L1* retrotransposon expression induces the interferon beta (*IFNβ*) inflammatory response (15, 16). This can lead to chronic inflammation (inflammaging), which is common in aged individuals and is associated with cellular senescence (16). In mammals, recognition of cytosolic RT DNA by *cGAS/STING* triggers the *NFκB* pathway and the *IFNβ* inflammatory response (14, 54). *Drosophila* does not have adaptive immunity, but many proteins that comprise the innate immune response are homologous to those that regulate the mammalian inflammatory response (54). In *Drosophila,* STING activates the *NFκB* homolog *Relish* to activate the immune defective (IMD) pathway and the expression of AMPs (54). This functions in parallel to the Toll pathway that acts through *Dorsal-related immunity factor (Dif),* another *NFκB* homolog, to activate the production of AMPs (55–57). We did not find the expression of components of the upstream components of the IMD and Toll pathways to be altered in *412* or *Roo* knockdown animals. However, several *Drosomycin* genes that encode AMPs downstream of the Toll pathway activation were downregulated upon TE knockdown. Lowering the expression of AMPs extends lifespan (58), suggesting that this may contribute to the lifespan changes seen in TE knockdown flies. While the mechanism by which *412* or *Roo* knockdown alters the expression of AMP genes remains to be determined, it may be linked to the upregulation of genes involved in proteolysis. In particular, genes of the *Jonah* serine hydrolase family are activated upon RNA viral infection and can influence the expression of *Drosomycins,* although the mechanism for this is unknown (47, 48). Jon proteins have a conserved function as their mammalian homolog, *Chymotrypsin like (CTRL),* is also involved in proteolysis within the gut (45, 59–62). Recently, a chymotrypsin/trypsin fusion protease has been utilized as a treatment for inflammation and to promote wound healing (63), suggesting a conserved link between these serine proteases and inflammation. We therefore hypothesize that the upregulation of the *Jonah* genes could dampen the immune system, and this could contribute to the extension of lifespan seen in animals with reduced expression of TEs.

Increased lifespan often coincides with improved stress resistance or other markers of health as illustrated by the insulin mutant and progeroid flies showing changes to oxidative stress and starvation and locomotion, respectively (37, 38). In contrast, knockdown of *412* or *Roo* increased lifespan without promoting observable stress resistance or delaying the onset of age-associated changes such as decreased mobility. Previous studies in *Drosophila* examining changes to heterochromatic and RNAi pathways to modulate TE activity did not examine these classic stress assays (11, 23). Thus, it is possible that TE expression-induced changes to lifespan occur without altering stress resistance, effectively uncoupling life- and healthspan. Defining the precise links between *412*, *Roo* and other TEs and their effect on cellular and organism function during aging will require additional genetic and molecular studies to elucidate the links between the *Jonah* genes, immunity, and aging.

## Materials and methods

### Fly strains

The following fly stocks were obtained from Bloomington Drosophila Stock Center: Act5C-Gal4 (BL #3954), *w^1118^* (BL#5905), UAS-dcas9-VPR (BL#67062). UH3-Gal4 was a gift from the Sparrow lab (Singh et al., 2014). UAS-shRNAs were generated according to the TRiP protocol using the pVALIUM20 vector (Addgene) (64). shRNA primers are listed in the primers table and were designed using the Designer of Small Interfering RNA webtool (65). UAS-shRNAs were inserted into the attP site at 86F (BL #24749) by BestGene. gRNA flies for CRISPRi were generated using the protocol from CRIPR fly design (66–72). The pCFD3 vector (Addgene) was used to insert a gRNA downstream of the U6:3 promoter, allowing for ubiquitous expression of the gRNA. gRNA transgenes were inserted into the attP site at 86F (BL #24749) by BestGene. UAS-dcas9:KRAB flies were created by In-Fusion® cloning system (Takara) using the pUAST-attB from the Drosophila Genomics Resource Center (# 1419) and dCas9-KRAB (Addgene SID4X-dCas9-KRAB; #106399). The primers are listed in the S8 Table and the PCR amplifications were performed using the CloneAmp™ HiFi PCR Premix (Takara). The pUAST-dCas9-KRAB plasmid was recombined into the attP site attP40 (BL #24749) by BestGene.

### Fly care

Fly food contained 80g malt extract, 65g cornmeal, 22g molasses, 18g yeast, 9g agar, 2.3g methyl para-benzoic acid and 6.35ml propionic acid per liter. Flies were kept at 25°C with a 12-hour light/dark cycle and 50% humidity. Day of eclosion is defined as day zero in all analyses. Adults were collected 2 days after the first flies eclosed, allowed to mate for a day, sorted by sex and allowed to age to the specified time point. Flies were kept at a density of 25 or less per vial. Flies were transferred to a new vial of food twice a week until they reached the desired age for the experiment. All of these studies used female flies.

### RNA-seq

Triplicate samples of 20 thoraces from adults were collected at day 20 from Act5C>shRNA (Control, 412#1, and Roo) and frozen at −80°C. RNA was extracted using TRIzol (Invitrogen) and sent to Novogene for quality assessment, library preparation, sequencing, and differential expression analysis. Sequencing was performed on using an Illumina platform with a sequencing by synthesis (SBS) method. HISAT2 was used to map the reads to the *Drosophila* genome (*dm6*), Novogene calculated the read counts and FPKM (fragments per kilobase of transcript per million base pairs sequenced) values and then used DESeq2 was used to perform differential expression analysis (73). DAVID was used to obtain gene ontology (GO) terms (42, 43). String was used to observe protein interactions of the enriched gene products (44). The data discussed in this publication have been deposited in NCBI’s Gene Expression Omnibus (74) and are accessible through GEO Series accession number GSE207160.

### qPCR

TRIzol was used to extract total RNA from 5 whole adult flies quantification of knockdown efficiency and heads and thoraces for examining expression of TEs in young and old *w^1118^.* RNA was DNase treated (Invitrogen) and cDNA was synthesized using the Verso cDNA kit (Thermo-Fisher AB1453A). 1-5μg of RNA was used for cDNA creation. PowerUp SYBR Green Master Mix was used to perform qPCR on the Applied Biosystems QuantStudio3 system. *rp49 (RpL32)* was used as the housekeeping gene to normalize relative gene expression changes. Experiments were performed in 3-5 biological replicates. An unpaired t-test with Bonferroni correction was used as the statistical test. Primers used in these experiments can be found in the S9 Table. (11, 75–82)

### Lifespan quantification

Adult flies were collected 48 hours after eclosion and mated for 24 hours. Flies were sorted by sex and genotype and placed at a density of no more than 25 flies per vial. The number of dead animals was counted twice a week and remaining live flies were transferred to new food vials. Lifespan experiments were performed in triplicate (separate crosses) and results pooled. A Dunnett test was used to compare survival curves. A Gehan-Breslow-Wilcoxon test was used to compare median lifespan. Difference in maximum lifespan was calculated by a permutation test (95th percentile) followed by two-sided t-test with correction for multiple comparisons.

### Oxidative stress survival

Flies were placed on 20mM paraquat (Sigma), 1% agar, 5% sucrose media at day 40 post eclosion. The number of dead animals were counted every 4-8 hours until all flies were dead. The experiment was performed in biological triplicate. A Dunnett test was used to compare survival curves.

### Starvation survival

Day 40 adult flies were provided with Whatman paper-soaked in water and the number of dead animals counted every 4 hours until all flies were dead. The experiment was performed in biological triplicate. A Dunnett test was used to compare survival curves.

### Endoplasmic reticulum stress survival

Flies were placed on 12 μM tunicamycin (Sigma), 1.5% agar, and 5% sucrose media at day 20 post eclosion. The number of dead animals were counted every day until all flies were dead. The experiment was performed in biological triplicate. A Dunnett test was used to compare survival curves.

### Negative geotaxis

Flies at days 5, 20, and 40 post eclosion were recorded while they were tapped down to the bottom of empty vials. After 10 seconds the number of flies in each third of each vial were counted. Flies were allowed a minute to recover before the experiment was repeated. The experiment was performed a total of three times in biological triplicate. A Chi-square test for trend was used to compare the locomotor activity of the different genotypes.

### Fecundity and Fertility

Females were mated with *w^1118^* males (1:1 ratio) at day 25 post eclosion at a density of approximately 20 flies per vial. Females were given 24 hours to lay eggs, then transferred to a new vial and the eggs per vial were counted every day for 5 days. Fecundity was calculated as the average number of eggs laid per vial per day. To assess fertility, the number of eggs laid was quantified and animals allowed to develop until eclosion at which point the number of adults flies were quantified. The fertility index was calculated by dividing the number of adult flies by the number of eggs laid per vial. A fertility index of 1 is defined as 100% fertile. A One-way ANOVA was used to calculate the differences in fecundity and fertility across genotypes.

### Thermal stress survival

To test sensitivity to cold stress, flies were placed in new food vials and placed at 4°C on ice for 15 hours at day 20 post eclosion. Flies were given 48 hours to recover at 25°C and the number dead flies counted. To test heat sensitivity, flies were placed in new food vials and placed at 37°C until about 80% of flies were immobile at the bottom of the vial with heat paralysis. Flies were given 48 hours to recover at 25°C and the number of dead flies were counted. Assays were performed in biological triplicate. A Fisher’s exact test was used to calculate the difference in survival across genotypes.

### Body weight and animal size

Zeiss Discovery.V12 SteREO with the AxioVision Release 4.8 software was used to capture pictures of adult flies using a ruler to show size. All images were processed using Adobe Photoshop. Flies were 2 days post eclosion for imaging and body mass quantification. To measure body mass, 10 flies of each genotype were placed in 1.5mL Eppendorf tube and weighed. This was done in biological triplicate. A One-way ANOVA was used to calculate body weight differences across genotypes.

### Purification and amplification of individual thoracic nuclei

40-60 thoraces from young (5 days old) or old (50 days old) *w^1118^* flies were dissected and single nuclei were prepared according to (83) with minor alterations. Briefly, instead of using a Polytron, thoraces were homogenized with an automatic pestle for 5 minutes on ice, transferred to a 7mL Dounce homogenizer and pulverized with 30 stokes of the pestle. Debris was filtered using a 20-micron filter and subsequently with a 10-micron filter (twice). Nuclei were then sorted into individual tubes using the CellRaft AIR® System. Genomic DNA from each nucleus was amplified by multiple displacement amplification and made into libraries according to (84). Libraries (21 per group) were subjected to paired end whole genome sequencing (WGS) on the Illumina 2500 platform at Novogene. The data discussed in this publication have been deposited in NCBI’s Sequence Read Archive and are accessible through BioProject accession number: PRJNA854389.

### Purification and genome amplification of IFM nuclei

UH3-Gal4/ +; +/+; UAS-Klar-KASH/+ flies were aged to either 5 (young) or 60 (old) days. 50 thoraces per group were dissected and single nuclei were prepared using the nuclei EZ isolation kit (Sigma) according to manufacturer instructions. Nuclei were then stained with DAPI, and fluorescence-activated cell sorting (FACS) sorted gaiting for DAPI (4’,6-diamidino-2-phenylindole) and GFP positive populations using the MoFloXDP at the Flow Cytometry Core Facility at Albert Einstein College of Medicine. The first gate selected for size by plotting forward scatter (FSC, size) against side scatter (SSC, granularity), with debris being outside the gate. 99.6% of the population was not debris. A log scale was used to visualize high signals from both axes in the same plot. The second gate selected for single nuclei by plotting SSC-width (doublets) against SSC-height (intensity). Single nuclei were 30.14% of the population. The last and final gate selected for intact IFM nuclei by plotting GFP (IFM) against DAPI (DNA). The nuclei were sorted into individual tubes and subject to single nucleus whole genome amplification (snWGA) using the REPLI-g® UltraFast Mini kit (Qiagen). The snWGA was performed by a multiple displacement amplification as in this previous publication (85). The snWGA amplicons were purified using AMPure XP magnetic beads (Agencourt). The snWGA amplicons were then subject to a locus drop out (LDO) test to screen for amplification of five regions dispersed throughout the genome, similar to previous studies (86). The Fast SYBR® Green Master Mix (Applied Biosystems) was used for the qPCR reaction. Nuclei with the most primer sets passing the LDO test were subjected to paired end sequencing (scWGS) on the Illumina 2500 platform at Novogene. Bulk/pooled unamplified genomic DNA from young abdominal segments was used as the control for genomic insertions already present in the strain. The data discussed in this publication have been deposited in NCBI’s Sequence Read Archive and are accessible through BioProject accession number: PRJNA854818.

### Bioinformatic analyses of single nucleus data

To control for genetic background germline TE insertions, genomic DNA was extracted from pooled young (day 5) thoraces from 50 flies and sequenced at Novogene. Fastq files were analyzed using the new pipeline Retrofind and also the published pipeline from the Bardin lab (27). Retrofind pre-filters input sequencing reads to require at least one mate pair to contain retrotransposon DNA sequences. Next, Retrofind conducts an alignment on the pre-filtered reads using the BWA mem aligner under strict conditions (87). Samtools coordinate sorts and Picard tools removes duplicates from the alignment (88). Reads inconsistent with a proper mate pairing or with larger than expected insert size are identified as a discordant read pair. Split reads are identified from reads aligning with soft or hard clipping above a threshold. Candidate split and discordant reads are aligned using BWA mem to a list of consensus sequences derived from Repbase (89). Candidate reads are then grouped into clusters using bedtools and designated as 5’ (left) and 3’ (right) junction reads (90). Next, a heuristic process is applied to the left and right junction to identify a retrotransposition that satisfies filtering options. If there is at least one right junction and one left junction split read, a target site duplication (TSD) prediction is made. There is an option to require a TSD prediction within a user-defined range. The default TSD size range we consider is 2-30 base pairs. The exported file includes an identification number that can be used to link reads of support to the transposition call. Lastly, Retrofind also outputs *de novo* assembly of supporting reads using the Megahit short read assembler (91) and a BWA mem alignment of assembled contigs to the genome. The reads of support and assembled contig alignment can be visually inspected using a genome browser. Retrofind was validated using the pipeline and methods described in (27).

*De novo* TE insertions detected in both young and old IFM nuclei compared to the bulk genomic DNA to exclude the insertions that are present within the germline. The genomic location of insertions identified in *w^1118^* were determined using ChIPSeeker (92). High quality insertions were defined as insertions detected within both pipelines and validated in IGV.

## Data Accessibility

RNA-seq data can be accessed through GEO Series accession number GSE207160. The thoracic WGS data can be accessed through the Sequence Read Archive (SRA) and are accessible through BioProject accession number: PRJNA854389. The indirect flight muscle WGS data can be accessed through the SRA BioProject accession number: PRJNA854818.

## Acknowledgements

We thank members of the Secombe, Vijg, and Baker labs for their insights throughout this project, particularly Shannon Lightcap who generated several transgenes used in this study, and Matanel Yheskel who helped with bioinformatic analyses. We are also very grateful to Alison Bardin and lab members for assistance with their TE insertion pipeline and Masako Suzuki, Jack Lenz and Kenny Ye for their genomics and genetics expertise. We are grateful for fly strains from the Bloomington *Drosophila* Stock Center (NIH P400D018537). This work was supported by NIH R01 AG053269 to J.S., T32AG023475 and T32GM007491 to B.K.S, the shared instrument grant 1S10OD023591-01, and the Einstein Cancer Center Support Grant P30 CA013330.

## Supporting information captions

**S1 Fig.**
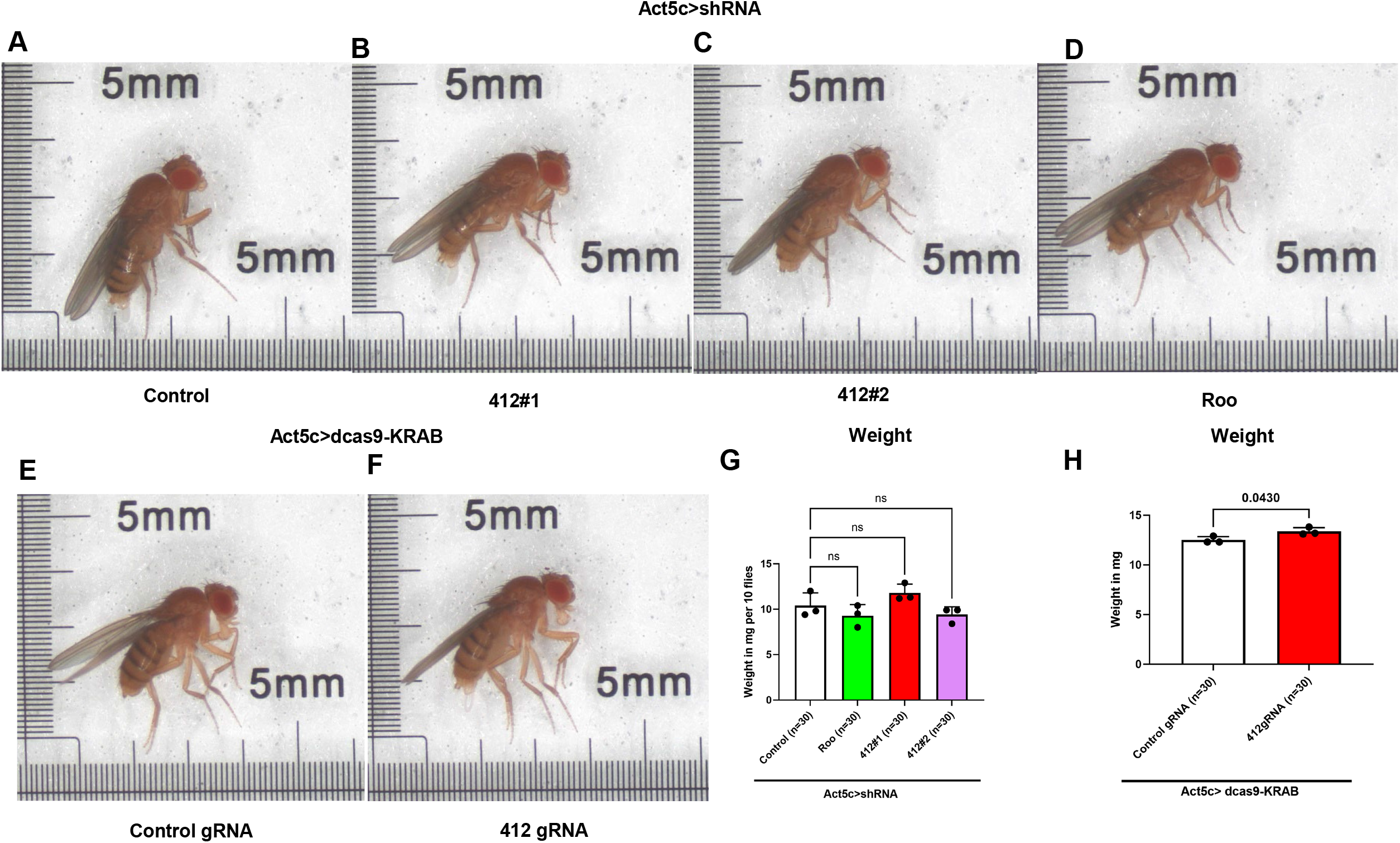
Morphology and body weight of *412* and *Roo* knockdown animals. (A) Picture of the morphology of an Act5c>control shRNA fly. (B) Picture of the morphology of an Act5c>412#1 shRNA fly. (C) Picture of the morphology of an Act5c>412#2 shRNA fly. (D) Picture of the morphology of an Act5c>Roo shRNA fly. (E) Picture of the morphology of a CRISPRi (Act5c>dcas9-KRAB) with control gRNA fly. (F) Picture of the morphology of a CRISPRi (Act5c>dcas9-KRAB) with 412gRNA fly. (G) Body weight of Act5c>shRNA flies in milligrams (mg) per 10 flies. Each dot represents a set of 10 flies. This experiment was done in triplicate. One-way ANOVA. ns (412#1) p= 0.3535. ns (412#2) p= 0.5904. ns (Roo) p= 0.5250. (H) Body weight of Act5c>dcas9-KRAB CRISPRi flies in milligrams (mg) per 10 flies. Each dot represents a set of 10 flies. This experiment was done in triplicate. Unpaired t-test. * p= 0.0430.

**S2 Fig.**
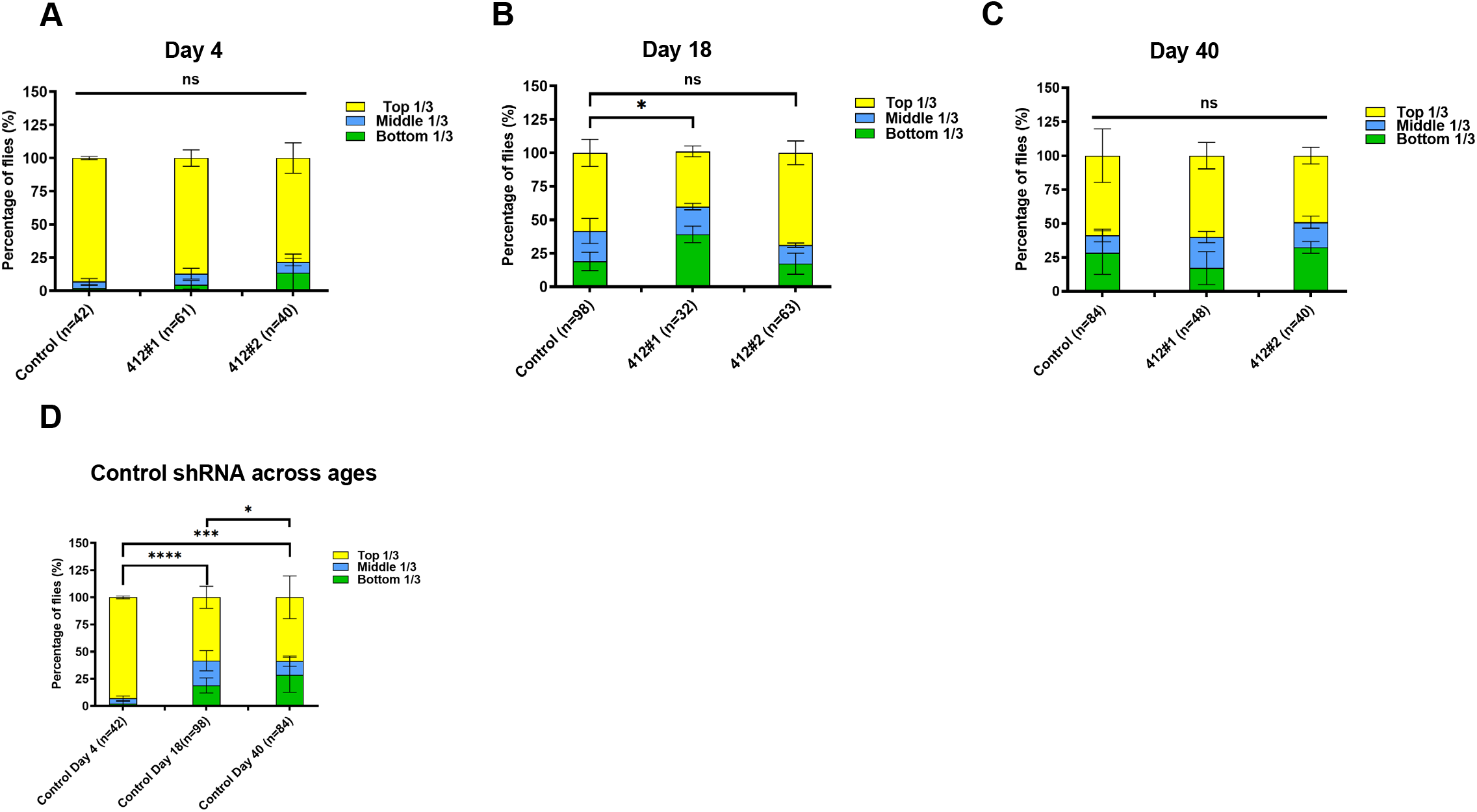
Negative geotaxis locomotion assay across age. (A) Act5c>shRNA day 4 measurement of locomotion via negative geotaxis assay. The percentages of flies in each third of the vial is displayed. Fisher’s exact test. ns (412#1) p= 0.5184. ns (412#2) p= 0.1119. (B) Act5c>shRNA day 18 measurement of locomotion via negative geotaxis assay. The percentages of flies in each third of the vial is displayed. Chi-square test for trend. * (412#1) p= 0.0332. ns (412#2) p= 0.4232. (C) Act5c>shRNA day 40 measurement of locomotion via negative geotaxis assay. The percentages of flies in each third of the vial is displayed. Chi-square test for trend. ns (412#1) p=0.4613. ns (412#2) p=0.3030. (D) Act5c>Control shRNA days 4, 20, and 40 measurement of locomotion via negative geotaxis assay. The percentages of flies in each third of the vial is displayed. Chi-square test for trend. **** (day 4 vs day 20) p<0.0001. *** (day 4 vs day 40) p= 0.0001. * (day 20 vs day 40) p= 0.0493.

